# Interactive visualization of whole eukaryote genome alignments using NCBI’s Comparative Genome Viewer (CGV)

**DOI:** 10.1101/2023.10.30.564672

**Authors:** Sanjida H Rangwala, Dmitry V Rudnev, Victor V Ananiev, Andrea Asztalos, Barrett Benica, Evgeny A Borodin, Nathan Bouk, Vladislav I Evgeniev, Vamsi K Kodali, Vadim Lotov, Eyal Mozes, Dong-Ha Oh, Marina V Omelchenko, Sofya Savkina, Ekaterina Sukharnikov, Joël Virothaisakun, Terence D. Murphy, Kim D Pruitt, Valerie A. Schneider

## Abstract

We report a new visualization tool for analysis of whole genome assembly-assembly alignments, the Comparative Genome Viewer (CGV) (https://ncbi.nlm.nih.gov/genome/cgv/). CGV visualizes pairwise same-species and cross-species alignments provided by NCBI using assembly alignment algorithms developed by us and others. Researchers can examine the alignments between the two assemblies using two alternate views: a chromosome ideogram-based view or a 2D genome dotplot. Whole genome alignment views expose large structural differences spanning chromosomes, such as inversions or translocations. Users can also navigate to regions of interest, where they can detect and analyze smaller-scale deletions and rearrangements within specific chromosome or gene regions. RefSeq or user-provided gene annotation is displayed in the ideogram view where available. CGV currently provides approximately 700 alignments from over 300 animal, plant, and fungal species. CGV and related NCBI viewers are undergoing active development to further meet needs of the research community in comparative genome visualization.

## Introduction

### Comparative genome visualization

Comparative genomics leverages shared evolutionary histories among different species to answer basic biological questions and understand the causes of disease. As sequencing costs have dropped and assembly algorithms have improved, there has been tremendous growth in the number of high-quality genome assemblies available in public archives, and the diversity in the organisms they represent. These data now make it possible to use comparative genomics approaches to explore more elements of biology and reveal the need for different types of analysis tools to support this exploration. The NIH Comparative Genomics Resource (CGR) maximizes the impact of eukaryotic research organisms and their genomic data to biomedical research (1). CGR facilitates reliable comparative genomics analyses for all eukaryotic organisms through community collaboration and a National Center for Biotechnology Information (NCBI) genomics toolkit. As part of CGR, we have created the Comparative Genome Viewer (CGV), a web-based visualization tool to facilitate comparative genomics research.

Graphical visualization of genomic data can illuminate relationships among different data types and highlight differences and anomalies; for instance, areas of a genome that are depleted in gene annotation, or have unusually high repeat content, or are more variable between species. Interactive genome browsers have become particularly valuable in recent years in helping biologists navigate large sequence datasets and more easily find genomic locations of interest to their specific research interest question. These visualizations can display molecular data that can help resolve competing hypotheses and also expose patterns that can spur additional research questions.

Linear genome browsers, such as the Genome Data Viewer (GDV) at NCBI (2), the UCSC Genomics Browser (3), and JBrowse (4), display molecular data as “tracks” laid out in parallel and anchored on a single genome assembly. Users can navigate to different chromosome regions and view gene annotation, repeats, and other types of data. Displays of known sequence variants (e.g., dbVar or dbSNP data) can aid in the analysis of gene or sequence level variation in a discrete genome region. While these browsers can be powerful tools to analyze many different types of data from a single genome concurrently, they are more limited in their ability to support comparative genome analysis. In particular, linear genome browsers cannot easily show whether a genome region that appears evolutionarily conserved has been rearranged (e.g., inverted or translocated) in one genome relative to another one, since the data is only displayed relative to a single genomic region in a single genome at a time.

Different types of two-dimensional visualizations have been proposed to facilitate analysis of larger scale genome structural differences between two or more genomes. These visuals include two-dimensional line graphs (also known as dotplots) (5, 6), circular diagrams (i.e., Circos plots) (7), and linear genome browsers that stack one assembly on top of another (8–10). Different types of visuals have advantages and disadvantages. Circos plots can show multiple datasets in one graphic but can be visually challenging to interpret and usually do not support views of sub-genomic regions. Dotplots can allow zooming to view chromosome or sub-chromosome regions but cannot easily or elegantly display gene or other annotation in the same visual. In order to better serve different research questions, some groups provide a choice of multiple different types of visuals for genome comparisons (11–13).

### Genome comparison data

Broadly, there are two types of whole genome comparison data that can be displayed in a comparative genome visualization tool. The first type of data is locations of gene orthologs. Orthology is typically determined using a protein homology-based method (e.g., BLASTP) in consideration with local gene order conservation (14, 15). This type of data can lend itself to straightforward “beads on string” visualizations that allows researchers to easily determine how syntenic gene regions have evolved across related species (15–17).

The second type of comparison data is whole genome assembly alignments (e.g., Mauve (18), LASTZ (19)), which are sequence-based and include both genic and intergenic regions. Whole genome alignments can be much more complex than simple gene ortholog locations but have the advantage of including alignments in regulatory regions and other regions not annotated as genes.

Here we introduce a new viewer tool at NCBI, the Comparative Genome Viewer (CGV), that is a key element of CGR. The main view of CGV takes the “stacked linear browser” approach — chromosomes from two assemblies are laid out horizontally with colored bands connecting regions of sequence alignment. Initial usability research with conceptual prototypes revealed that this type of visual was the easiest to interpret for scientists from a broad range of research expertise in genomics. We display whole genome pairwise assembly-assembly alignments in CGV. These sequence-based alignments can be used to analyze gene synteny conservation but can also expose similarities in regions outside known genes e.g., ultraconserved regions that may be involved in gene regulation. Because CGV is a web-based application, researchers do not have to install or configure software or generate their own comparison files before they can begin using it for their research. Below we describe some of the features of CGV and provide examples of how visualization in this tool can generate insights into genome structure and evolution.

## Results

### Overview of CGV

We developed a web application, the Comparative Genome Viewer (CGV) (https://ncbi.nlm.nih.gov/genome/cgv/), to aid in comparing genome structures between two eukaryotic assemblies. CGV facilitates analyses of genome variation and evolution between different strains or species or strains, as well as evaluation of assembly quality between older and newer assemblies from same species.

Alignments are generated at NCBI using BLAST (20) or LASTZ-based algorithms (19), or imported from the UCSC Genomics Institute (https://hgdownload.soe.ucsc.edu/downloads.html) or from other research groups (e.g., HPRC, https://humanpangenome.org/). Shorter alignments are merged where possible; however, because of repeats and gaps in the alignments, even very similar chromosomes may be broken down into many alignment segments. Refer to *Materials and Methods* for more detail on how we generate whole-genome assembly alignments and load them into the viewer.

The CGV home page provides a menu where users can select from available species and assembly combinations (Fig 1A). We frequently add new alignments as high-profile assemblies become available and in response to requests from the scientific community. As of October 2023, we provided a selection of almost 700 alignments from over 300 eukaryotic species (Fig 1B).

**Fig 1.**
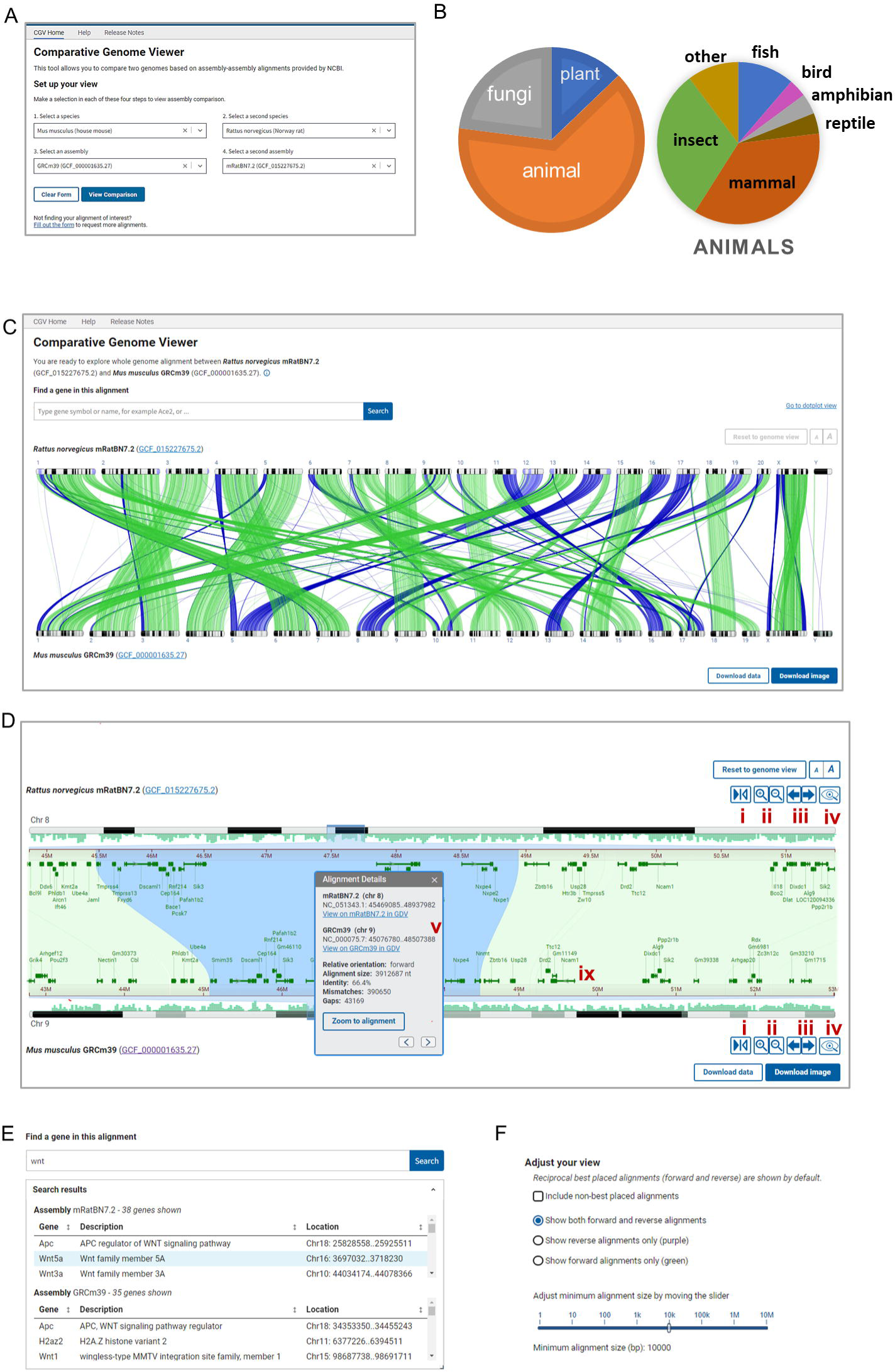
Overview of CGV. (A) CGV selection menu. (B) Taxonomic distribution of species represented by alignments in CGV. (C) CGV ideogram view of whole genome assembly alignment. Buttons in the lower right provide download access for complete alignment data or an SVG image of the current alignment view. https://www.ncbi.nlm.nih.gov/genome/cgv/browse/GCF_015227675.2/GCF_000001635.27/27835/10116 (D) CGV zoomed to a chromosome-by-chromosome view, with information box shown. i. flip orientation ii. zoom in/out iii. pan left/right iv. view assembly in Genome Data Viewer (GDV) v. information panel. https://www.ncbi.nlm.nih.gov/genome/cgv/browse/GCF_015227675.2/GCF_000001635.27/27835/10116#NC_051343.1:44657546-50471623/NC_000075.7:41365809-52285446/size=10000 (E) CGV search interface and sample search results. (F) Filtering and configure options for ideogram view.

CGV’s main view (the “ideogram view”) displays pairwise alignments as colored connectors linking the chromosomes in the two assemblies (Fig 1C). The view is filtered by default to show only reciprocal best hits between assemblies to facilitate the analysis of orthologous genomic regions. The researcher can choose to show the non-best placed alignments to expose close sequence duplications or homologues (Fig 1F). Users of CGV can also filter alignments in view by size (e.g., to only show large alignments blocks) or by orientation (e.g., to only show regions that have undergone a potential inversion). The complete alignment data in GFF3 and human-readable formats like XLSX can be downloaded from the viewer for a researcher’s own use.

Users can select a chromosome from each assembly to zoom to the alignments for this chromosome. They can navigate further within this chromosome comparison using the zoom in/out and pan buttons or by using the mouse to pinch-zoom or drag to pan. Users can zoom directly to a particular region of a chromosome by dragging their cursor over the coordinate ruler or the ideogram for either assembly. Double-clicking on a selected alignment segment will synchronously zoom both the top and bottom assembly on the aligned coordinates, so that they are stacked on top of one another (Fig 1D).

Where available, RefSeq or assembly-submitter provided gene annotation is displayed on the chromosomes (Fig 1D). Similarities in gene order denote regions of synteny, while discrepancies can point to evolutionarily or biologically significant differences, or assembly errors, particularly if evaluating different assemblies from the same species. Researchers can use the search feature in CGV to find their gene of interest by name or keyword, and subsequently navigate to the location of the gene in the viewer (Fig 1E). If the gene region is aligned, the viewer will simultaneously navigate to the aligned location, which may contain the gene’s known or putative ortholog on the second assembly. The “flip” button allows the user to reverse one chromosome to see inverted alignments displayed in the same relative orientation, which may aid in the detection of discrepancies in gene annotation in regions that are locally syntenic between the two assemblies. Once a user has completed their analysis of a region of interest, they can export the image as an SVG to adapt for use in publications and presentations.

We provide additional information for each alignment segment in a pop-over panel (Fig 1D). This panel reports the chromosome scaffold accession and sequence coordinates of the alignment on each assembly, as well as the percent identity, number of gaps and mismatches, and alignment length. While the ideogram view in CGV does not display specific nucleotide bases, users can open another panel from the right-click menu that shows the alignment sequence. They can also download the alignment FASTA file of a particular alignment segment for downstream analysis, such as BLAST search or primer design. Moreover, researchers can also navigate from CGV to NCBI’s genome browser, the Genome Data Viewer (GDV) (2). GDV can display the assembly-alignment data viewed in CGV as a linear track alongside additional data mapped onto a genome assembly, such as detailed transcript and CDS annotation, repeats, GC content, variation data, or user-provided annotations. Zooming to a location within GDV can reveal granular differences in nucleotide sequence or gene exon or CDS annotation between the two assemblies.

In addition to the main ideogram-based view, the Comparative Genome Viewer also provides a two-dimensional dotplot view of the pairwise genome alignment (Fig 2A). The dotplot shows aligned sequence locations in one assembly on the X-axis plotted against aligned locations on the second assembly on the Y-axis. Alignments in the reverse orientation are plotted with an opposite slope and in a different color (purple) than alignments in the same orientation (green), making it easier to identify inversions and inverted translocations. The CGV dotplot shows both reciprocal best-placed and non-best placed alignments. As a result, compared to the ideogram view, this plot may more easily expose differences in copy number between two assemblies, such as segmental duplications or differences in genome or chromosome ploidy. Users can select a pair of chromosomes in the whole genome plot (i.e., a “cell” in the plot) and zoom into them on a full screen, where smaller alignment segments may be more easily interpretable (Fig 2B). Once a researcher has discovered a chromosome pair of interest, they can navigate back to the ideogram view to conduct even more fine-grained analysis, including examining gene annotation and observing very short alignment segments that were beyond the resolution of the dotplot.

**Fig 2.**
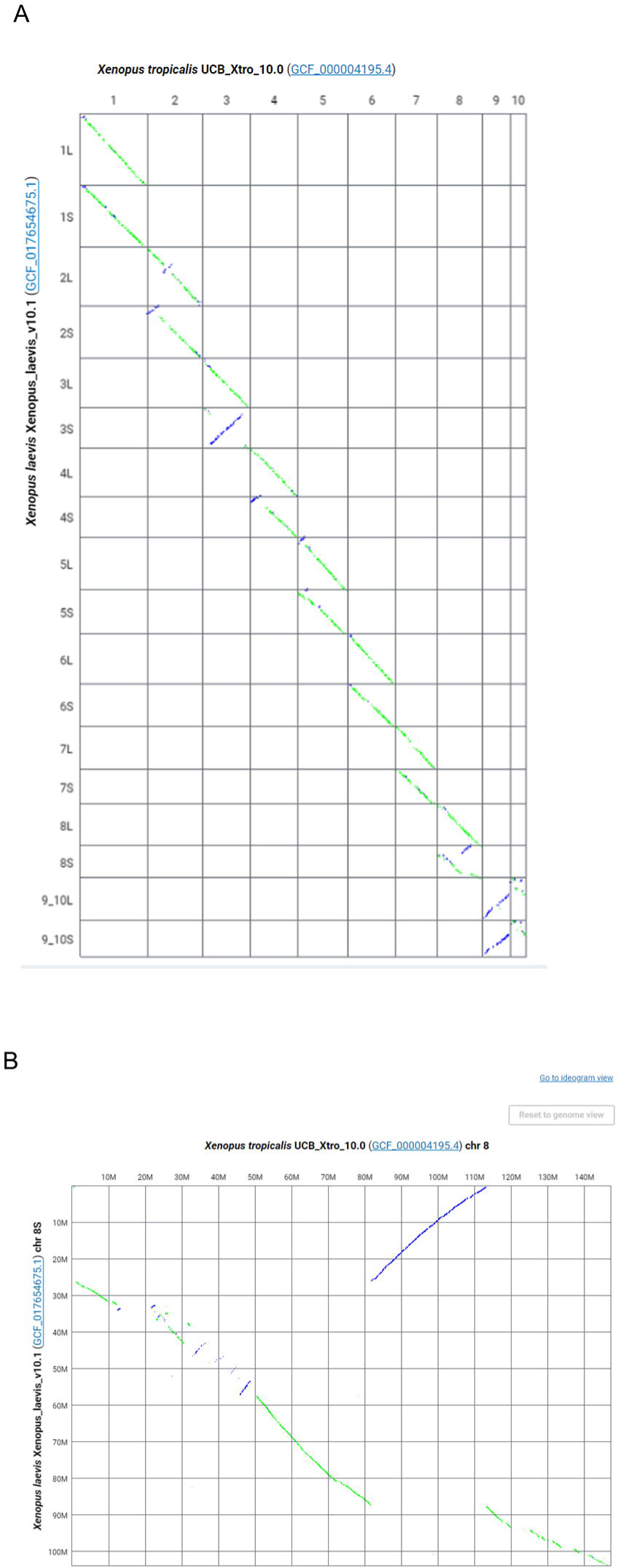
CGV dotplot view of Xenopus laevis and Xenopus tropicalis alignment. https://ncbi.nlm.nih.gov/genome/cgv/plot/GCF_000004195.4/GCF_017654675.1/38475/8355 (A) Full genome dotplot. (B) Dotplot of chromosome 8 of Xenopus tropicalis vs chromosome 8S of Xenopus laevis.

*Analysis using CGV: Conservation of linkage groups with local rearrangement of synteny* CGV can aid in detecting unusual patterns in genome evolution in different taxa. Researchers had previously observed that genomes from Drosophila species conserve gene content within linkage groups, known as Muller elements, corresponding to chromosomes or large sub-chromosome regions. Within these Muller elements, gene order can be reshuffled extensively in one species relative to another (21). For alignment between *D albomicans* vs *D melanogaster*, the CGV ideogram view shows restriction of alignment from each chromosome in one genome to a particular chromosome or chromosome region (i.e., linkage group) in the other genome (Fig 3A). However, within a chromosome-chromosome pair, sequence alignment is broken into many small fragments whose relative order is not conserved. This fragmentation of alignment is more clearly visible in the CGV dotplot, which shows that pairwise alignments are restricted to a single chromosome pair, but appear in a scattered pattern, suggesting that the sequence and gene order has been extensively rearranged within chromosomes (Fig 3B).

**Fig 3.**
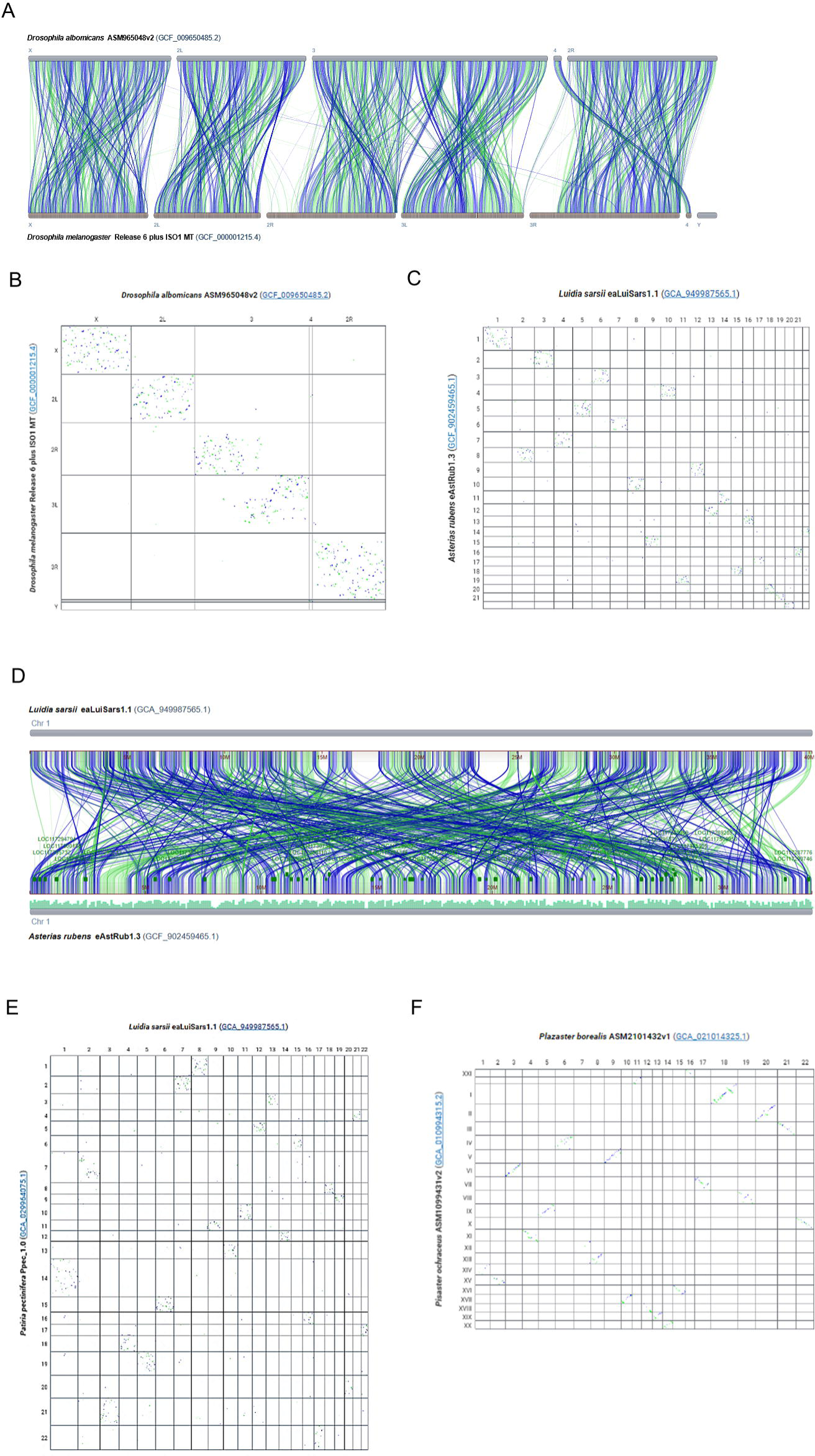
CGV shows conservation of linkage groups in the absence of conservation of gene order. (A) CGV ideogram view of alignment between Drosophila albomicans and Drosophila melanogaster genomes. Alignments are restricted to a single chromosome or chromosome region. https://ncbi.nlm.nih.gov/genome/cgv/browse/GCF_009650485.2/GCF_000001215.4/40865/0 (B)CGV dotplot view of alignment between Drosophila albomicans and Drosophila melanogaster demonstrates that sequence order is “scrambled” within linkage groups, as demonstrated by a scatter pattern indicating many short rearranged alignments. https://www.ncbi.nlm.nih.gov/genome/cgv/plot/GCF_009650485.2/GCF_000001215.4/40865/0 (C) CGV dotplot view of alignment between starfish species Luida sarsii and Asteria rubens with similar scatter pattern to Drosophila alignments. https://www.ncbi.nlm.nih.gov/genome/cgv/plot/GCA_949987565.1/GCF_902459465.1/41045/0 (D) CGV ideogram view of alignment between chromosome 1 of Luida sarsii and chromosome 1 of Asteria rubens. These chromosomes align to each other across their length, but the alignment is broken into multiple short segments which are extensively rearranged. https://www.ncbi.nlm.nih.gov/genome/cgv/browse/GCA_949987565.1/GCF_902459465.1/41045/0#OX465101.1/NC_047062.1/size=1,firstpass=0 (E) CGV dotplot view of alignment between starfish species Luida sarsii and Patiria pectinifera with similar scatter pattern to Drosophila alignments. https://www.ncbi.nlm.nih.gov/genome/cgv/plot/GCA_949987565.1/GCA_029964075.1/41165/0 (F) CGV dotplot view of alignment between starfish species Plazaster borealis and Pisaster ochraceus. Alignments show less scatter and more of a diagonal slope, indicating more conservation of sequence order between these two species’ genomes. https://www.ncbi.nlm.nih.gov/genome/cgv/plot/GCA_021014325.1/GCA_010994315.2/41175/466999

We observed a similar pattern when comparing some genomes from different starfish species using CGV. When looking at pairwise alignments between *Asterias rubens*, *Patiria pectinifera*, and *Luida sarsii* species, sequences from a chromosome from one genome mainly align to a single other chromosome in the other species. However, within a pairwise chromosome alignment, the sequence order is rearranged, resulting in a scatter pattern in the dotplot (Fig 3C, E). The ideogram view can show the alignment fragmentation and rearrangement in more granular detail (Fig 3D). We also noted that some starfish pairs show more conservation of location synteny (22) (Fig 3F), consistent with measured sequence distance based on shared k-mers (Mash distance is 0.128 compared to other pairs with a Mash distance >0.3).

Bhutkar et al (21) speculated that the need to keep certain genes in the same regulatory environment may result in conservation of genes within linkage groups, even in the absence of selective pressure to maintain the gene order. More recently, conservation of macrosynteny with extensive small-scale sequence rearrangement was also detected in comparisons between other invertebrate species, such as cephalopods, cnidarians, jellies, and sponges (16, 23, 24). These rearrangements were used to parse the phylogenetic relationships within this clade. We demonstrate here that CGV can aid researchers in detecting and analyzing this phenomenon in starfish and other evolutionarily varied taxa.

### Analysis using CGV: Detection of amylase family expansion

CGV can uncover potential copy number differences in segmental gene families. These differences may appear as gaps in the alignment in otherwise syntenic gene regions. Segmental insertions or deletions may be too small to be apparent on the whole genome or whole chromosome alignment but can be detected when searching and navigating to a gene of interest.

Initial analysis of the complete human telomere-to-telomere CHM13 genome revealed an expansion of amylase genes on chromosome 1 compared to the GRCh38 reference assembly (25). This expansion can be validated in CGV: a search for ‘amylase’ in the alignment between GRCh38 and T2T-CHM13v2 assembly finds six matches to this gene name in the GRCh38 and twelve in the CHM13 assembly (Fig 4A). Zooming out in the region of the *AMY1A* gene on chromosome 1 reveals a nearby sequence segment in the CHM13 assembly that is not aligned to GRCh38 (Fig 4B). This region contains numerous annotated loci that lack official nomenclature (i.e., LOC); six of these loci are described as ‘alpha-amylase’. Therefore, there are at least six additional alpha-amylase genes in CHM13 genome compared to the GRC reference. It is possible that copy number of this gene is variable in humans; it is also possible that the GRC reference genome represents fewer than the typical number of gene copies. While these two human assemblies are likely to be high quality, other assemblies in other species may be of lower quality, and differences in copy number observed in CGV may reflect assembly or annotation errors.

**Fig 4.**
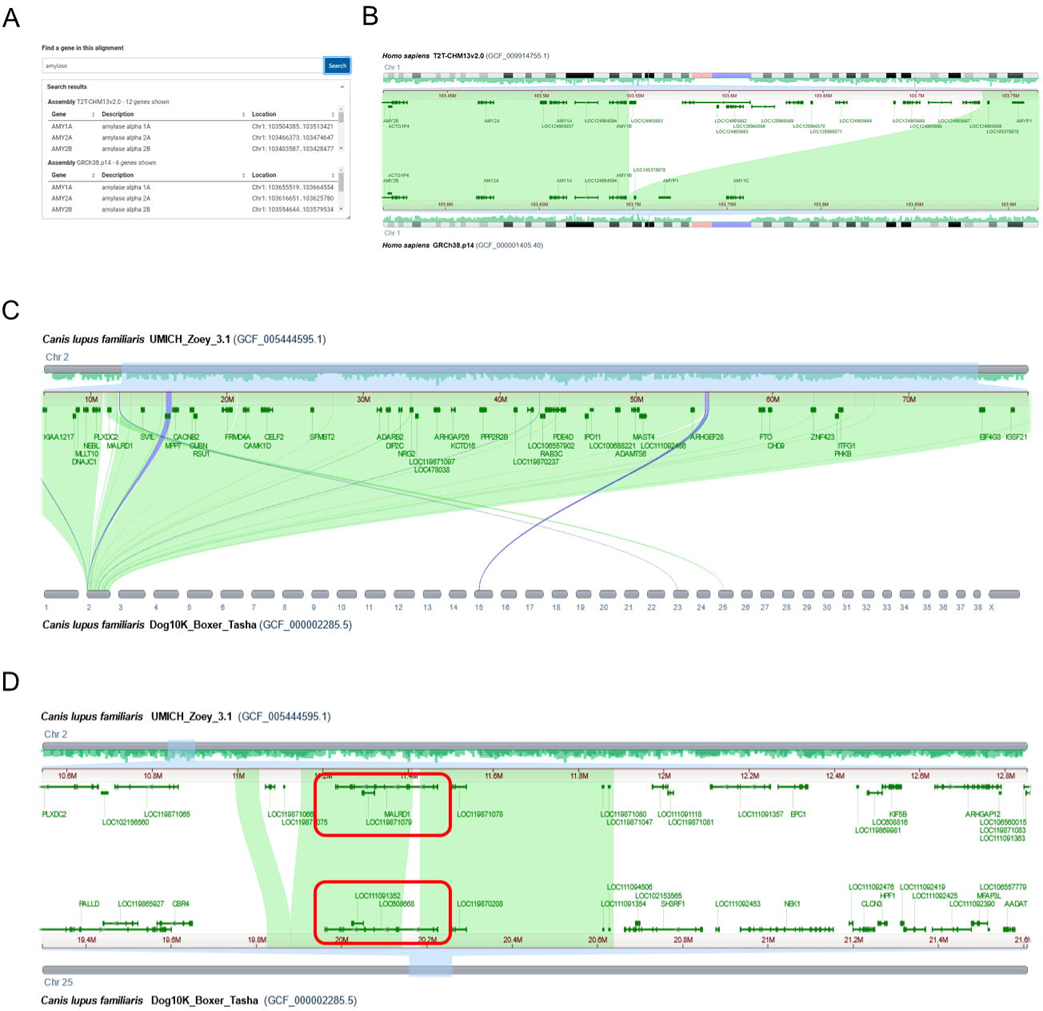
CGV can help uncover gene duplications and rearrangements in closely related genomes. (A) Gene search of an alignment between two human assemblies in CGV finds twelve amylase gene family members in the human T2T-CHM113v2.0 assembly and six amylase gene family members in GRCh38.p14. (B) CGV view showing that T2T-CHM13v2.0 contains an insertion relative to GRCh38.p14, which appears as an unaligned region on chromosome 1. This insertion contains additional alpha-amylase family members. A popup label (tooltip) indicates one of these additional family members. https://www.ncbi.nlm.nih.gov/genome/cgv/browse/GCF_009914755.1/GCF_000001405.40/23025/9606#NC_060925.1:103415704-103764412/NC_000001.11:103566852-103915505/size=1000,firstpass=0 (C) CGV view showing that chromosome 2 of Canis lupus familiaris (dog) UMICH_Zoey_3.1 align to chromosomes 2, 15, 23, and 25 of Dog10K_Boxer_Tasha. https://ncbi.nlm.nih.gov/genome/cgv/browse/GCF_005444595.1/GCF_000002285.5/17685/9615#NC_049262.1:6542815-78714085//size=10000 (D) UMICH_Zoey_3.1 assembly chromosome 2 alignment to Dog10K_Boxer_Tasha chromosome 25 contains the MALRD1 gene, which is annotated as LOC608668 in the Tasha assembly (boxed in red). Gene synteny is not conserved outside of the region of assembly-assembly alignment.

### Analysis using CGV: Possible gene translocation between two dog assemblies

For closely related strains or species, CGV can help uncover and validate structural anomalies, such as where gene order synteny has been disrupted. Visual inspection of the whole genome CGV ideogram view of alignment between two dog genomes – the Great Dane Zoey (UMICH_Zoey_3.1) and the boxer Tasha (Dog10K_Boxer_Tasha) – indicated a region that aligned to chromosome 2 in the Zoey assembly and chromosome 25 in the Tasha assembly (Fig 4C). This region contains the *MALRD1* gene in the Zoey assembly, which is shown to align to a *MALRD1* homolog annotated as *LOC608668* in Tasha (Fig 4D). CGV alignments indicate that *LOC608668* is likely the Tasha *MALRD1* gene; there are no better alignments to *MALRD1* detected in CGV or by an independent BLAST search of the Tasha genome.

Zooming out in the aligned region of *MALRD1* indicates that gene synteny is not conserved outside of the *MALRD1* gene region (Fig 4D). It appears that this gene has been rearranged from chromosome 2 on the Zoey assembly to chromosome 25 on Tasha. It is also possible that the Tasha genome may have been misassembled in this region, and the *MALRD1* gene sequence is properly situated on chromosome 2 within the otherwise conserved syntenic block. A researcher would need to further examine the quality of the Tasha assembly in this region to distinguish these possibilities, for instance, by examining the sequencing reads for the Tasha assembly or viewing a HiC map of assembly structure.

If this anomaly represents a true difference between the two genomes, it could prove biologically significant. The translocation of *MALRD1* may have placed it in a different gene regulatory environment in the boxer (Tasha) genome, which could possibly result in different levels or patterns of gene expression. The human ortholog of *MALRD1* was shown to be involved in bile acid metabolism (26). This gene region was also genetically linked to Alzheimer’s disease (27). Therefore, if valid and not assembly artifacts, differences like these could provide insight into human health.

## Discussion

We describe here a new visualization tool for eukaryotic assembly-assembly alignments, the Comparative Genome Viewer (CGV). We developed this web application with a view toward serving both expert genome scientists as well as organismal biologists, students, and educators. Users of CGV do not have to generate their own alignments or configure the software using command line tools. Instead, they can select from our menu of available alignments, access a view immediately in a web application, and start their analysis. We are continuing to add new alignments regularly and invite researchers to contact us if assemblies or organisms of interest are missing. We continue to do periodic outreach to the community to help us improve our visual interfaces so that they are simple, intuitive, and accessible.

CGV displays whole genome sequence alignments provided by NCBI; users cannot currently upload their own alignment data or choose assemblies to align in real time. There are both technical and scientific considerations to allowing researchers to select and align assemblies themselves. Currently, whole genome assembly-assembly alignments take several hours to days, using up to one thousand CPU processing hours per pairwise alignment of larger genomes, such as those for mammalian or plant assemblies. Moreover, whole genome alignments are difficult to generate past a certain genetic distance (i.e., Mash > 0.3). Not only is alignment between distantly related genomes computationally expensive, but the alignments themselves may be of limited research value. These alignments will likely have sparse and short segments that may correspond only to the most highly conserved coding sequence (CDS) (Fig 5; S1 Table). We suggest protein similarity or gene orthology-based alignments as more appropriate for comparisons between distantly related organisms. At present, we manually vet requested alignments to make sure that assemblies are complete, of high assembly quality (e.g. high scaffold N50 or BUSCO scores (28)), and a reasonable evolutionary distance. This review ensures that alignments will be useful to both the original requester and others in the research community.

**Fig 5.**
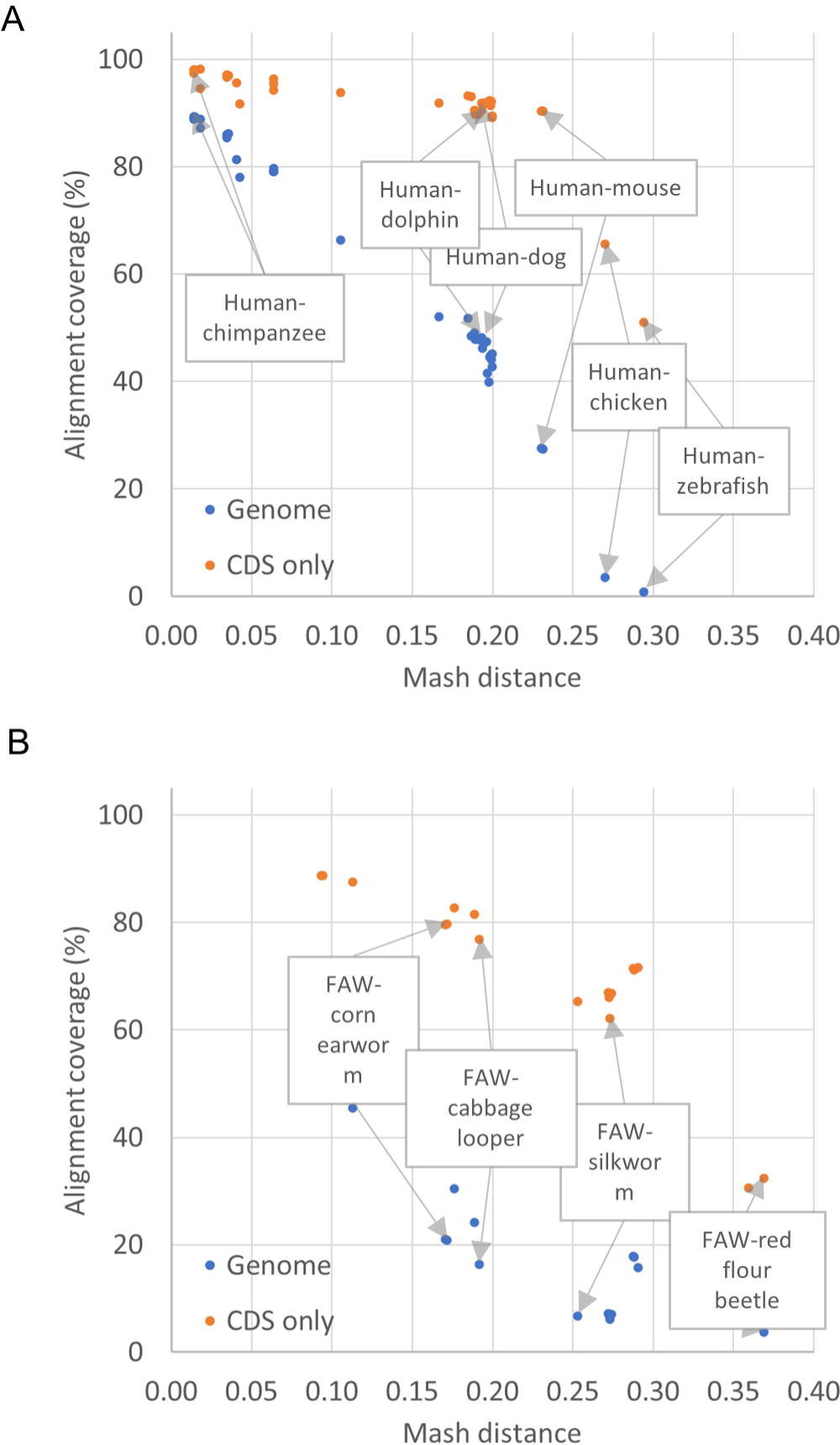
Genome and CDS coverage of assembly-assembly alignments relative to Mash distance. Percentages of total target genome or CDS nucleotides covered by the ungapped alignments are plotted against the Mash distance between the pair of genomes. (A) Alignments between the human GRCh38.p14 assembly and other vertebrates. (B) Alignments of fall army worm (FAW, Spodoptera frugiperda, a major insect agricultural pest) and related insects. At lower Mash distances, the whole genome alignments cover most of the genome and CDS. At Mash distances greater than 0.25 or 0.3, the alignment covers less than 20% of the genome overall, and between 30% and 60% of the CDS. Refer to S1 Table for the data used to populate these graphs.

Many research questions in comparative biology may be best answered by simultaneously visualizing alignments among more than two assemblies. We are exploring user needs and existing tools when it comes to multigenome alignment visualization, such as visualization of pangenome data. Our lessons from developing CGV will prove valuable in this upcoming initiative.

## Materials and Methods

### Preparation of assembly-assembly alignments

Genome assemblies are aligned using a two-phase pipeline first described in Steinberg et al (29), with adaptations for cross-species alignments. In the first phase, initial alignments are generated using BLAST (20) or LASTZ (19), or imported from a third-party source such as UCSC (30). In the second phase, alignments are merged and ranked to distinguish reciprocal-best alignments from additional alignments that are locally best on one assembly but not the other. The resulting alignment set is omnidirectional and can be used to project information from query to subject assembly or vice versa.

In the first phase, for both BLAST and LASTZ, repetitive sequences present in the query and target assemblies are soft-masked using WindowMasker (31). The default parameters are usually suitable for aligning assembly pairs within the same species. However, more aggressive masking is required when aligning cross-species assemblies. The masking rate is adjusted with the parameter `t_thres_pct` set to 99.5 (default, for BLAST same-species), 98.5 (for BLAST cross-species), or 97.5 (LASTZ cross-species), or lower for some genomes with extensive and diverse repeat composition. The 97.5 - 98.5 values typically result in a masking percentage similar to RepeatMasker (http://www.repeatmasker.org) with a species-specific repeat library, with the advantage of not needing to define repeat models beforehand.

Genome assemblies are aligned using either BLAST or LASTZ. The selection of the aligner and specific parameters depends on the level of similarity between the assemblies. We use Mash (32) to compute the approximate distance between two assemblies (Fig 5). BLAST is employed for aligning pairs of assemblies belonging to the same species, as well as cross-species assembly pairs with a Mash distance of less than 0.1. An exemplar BLAST command is: `blastn -evalue 0.0001 -gapextend 1 -gapopen 2 -max_target_seqs` 250 -`soft_masking true -task megablast -window_size 150 - word_size 28` A BLAST `word_size` of 28 is used for pairs of assemblies with Mash distances below 0.05, such as human and orangutan, while a `word_size` of 16 is used to enhance sensitivity for more distant cross-species pairs with Mash distances ranging from 0.05 to 0.1, such as human and rhesus macaque.

Assembly pairs with Mash distances exceeding 0.1, such as human and mouse assemblies, are aligned using LASTZ. The `make_lastz_chains` pipeline (19) is employed to generate alignments between query and target assemblies in UCSC chain format. The default parameters are often adequate to produce satisfactory alignments for many assembly pairs, though some distant assembly pairs (e.g., Mexican tetra-medaka) warrant changes such as the use of a different substitution matrix (`BLASTZ_Q=HoxD55`).l’.

Alignments generated with the `make_lastz_chains` pipeline or precomputed alignments imported from UCSC are in chain format. The UCSC chainNet pipeline (33) is run on query x target and target x query alignment sets separately so that the alignments are ’flattened’ in a way that the reference sequences are covered only once by the alignments in each set, and the two chainNet outputs are concatenated.

In the second phase, alignments are converted to NCBI ASN.1 format and processed further for the NCBI’s CGV and GDV browsers. In this phase, the full set of BLAST or LASTZ-derived alignments are processed to merge neighboring alignments and split and rank overlapping alignments to identify a set of best alignments (S1A Figure). Merging is accomplished on a sequence-pair-by-sequence-pair basis, and ranking is accomplished globally for the assembly pair. The process is designed to find a dominant diagonal among a set of potentially conflicting alignments.

Merging involves the following steps. First, when applicable, alignments based on common underlying sequence components of the assemblies (e.g., the same BAC component used in both human GRCh37 and GRCh38) are identified and merged into the longest and most consistent stretches possible. Second, adjacent alignments are merged if there are no conflicting alignments. Third, alignments are split on gaps using a default threshold of 50 bp (for the same or closer species) or 50 kb (for more distant species), or longer than 5% of the alignment length. Alignments are also split at any point where they intersect with overlapping alignments (S1A Figure). Duplicate or low-quality alignments fully contained within higher-quality alignments are dropped.

After merging and splitting, alignments are subsequently processed using a sorting and ranking algorithm (S1B Figure). Alignments are sorted based on a series of properties, including the use of common components, assembly level (alignment to chromosomes preferred over alignment to unplaced scaffolds), total sequence identity, and alignment length. The alignments are then scanned twice, once each on the query and subject sequence ranges, to sort out reciprocal best-placed (also referred to as “first pass” or reciprocity=3) and non-best placed (also referred to as “second pass” or reciprocity=1 or 2) alignment sets. Finally, all alignments in each reciprocity are merged again to stitch together adjacent alignments with no conflicting alignments into the longest representative stretches.

Assembly-assembly alignments are stored in an internal database available for rendering in NCBI’s CGV and GDV browsers. The alignment data are publicly available in GFF3 and ASN.1 formats at https://ftp.ncbi.nlm.nih.gov/pub/remap/.

For display in CGV, assembly alignment batches are filtered to keep only alignments that contain chromosome scaffolds as both anchors and targets, since non-chromosomal scaffolds are not displayed in this viewer. Alignments are converted into a compact binary format designed to keep only the synteny data required for display. This preparation step is done by programs written in C++ and bash scripts that tie them together.

### Technical architecture of CGV

CGV operates on a two-tier model, with a front end implemented using HTML/JavaScript running in the user’s web browser and a backend running at NCBI. The graphical rendering is done on the front end using modern WebGL technologies. The main advantage of this approach is speed and fluidity of the user interface since most of the alignment data needed to be rendered is sent to the front end at the initial load and there are no additional roundtrips to the server when the user interacts with the page (e.g., panning or zooming). Using front end graphical rendering also reduces the network traffic between NCBI and the end user, which makes CGV more responsive.

The backend of the CGV application resolves internal alignment identifiers to an alignment data file that the front end can use for generating graphical images. The back end is implemented as an industry-standard gRPC service written in C++ and running in a scalable NCBI service mesh (linkerd, namerd, consul). When a CGV view is initially loaded, our gRPC service requests the alignment data needed for the particular page. On graphical pages, gRPC resolves an alignment identifier to a URL with prepared synteny/alignment data and the page loads the file at this URL into a WASM module which is written in C++ and compiled with Emscripten. The WASM module serves this data on demand to the page’s JavaScript code which uses it for building the image and all user interactions.

In parallel, the list of assembly-assembly alignments and their metadata is sent to the selection menu (i.e., “Set up your view”) on the CGV landing page. This allows the selection menu to report scientific and common species names, assembly accessions, and assembly names.

The alignment selection menu on the CGV landing page is a traditional web form page. We utilize the NCBI version of United States Web Design System (USWDS) design standards and components (https://www.ncbi.nlm.nih.gov/style-guide) in order to unify graphical design with other US government pages.

On the more graphically intensive ideogram and dotplot pages, graphical rendering is done in the user’s web browser using WebGL using d3.js or pixi.js libraries, which allows for efficient interactivity, scalability, and fluidity of user interaction. Other elements of the page use jQuery/extJS and USWDS components. CGV reuses chromosome ideograms initially developed for NCBI’s Genome Data Viewer (2).

Gene annotations shown in CGV are obtained from NCBI’s public databases using NCBI Entrez Programming Utilities (E-utilities) (https://www.ncbi.nlm.nih.gov/books/NBK25501/).

Annotation-build specific gene search is provided by an NCBI Datasets (https://www.ncbi.nlm.nih.gov/datasets/) gRPC service.

### Design of CGV Application

A philosophy of user-centered design, which puts user needs at the forefront of decision- making, was an integral element in the development of the Comparative Genome Viewer (CGV). Participants for user research were recruited from members of the genomics research community who provided their contact information through feedback links on the CGV application and other sequence analysis tools at NCBI. Some of these researchers were previously familiar with CGV, while others had little or no experience with this tool. Data from user research testing sessions was compiled and analyzed for patterns in behavior, thereby allowing the team to validate that the design was moving in a direction that facilitated analysis of sequence alignment data. To date, we have conducted user research with over 30 different experts in the field of comparative genomics. We also evaluate the application for Section 508 compliance, which also helps insures CGV performs well on mobile devices and is accessible to users with limited or no visibility.

## Acknowledgments

We thank Anne Ketter and Emily W Davis for help with project planning and coordination. We thank Guangfeng Song for aid with user experience research and analysis. Thanks to Wayne Matten, Michelle Formica-Frizzi, and others in the NCBI customer service team for marketing, webinars, and video tutorials. Anatoliy Kuznetsov and Andrei Shkeda conducted early technical design work that influenced the architecture of the CGV application. Deanna M. Church and Mike DiCuccio participated in initial design and testing of the assembly alignment protocol. Finally, we thank members of the greater genomics research community who have participated in usability sessions and provided feedback and alignment requests to CGV. CGV was developed as a part of the National Institutes of Health’s Comparative Genomics Resource (CGR) (https://www.ncbi.nlm.nih.gov/comparative-genomics-resource/). This work was supported by the National Center for Biotechnology Information of the National Library of Medicine (NLM), National Institutes of Health.

---

S1 Table. Alignment coverage at different Mash distances for selected assembly pairs.

S1 Figure. Merging, sorting, and ranking assembly-assembly alignments. (A) Flowchart showing that adjacent alignment segments are merged. Subsequently, alignments are split once again at large gaps. (B) Flowchart showing how overlapping alignments are separated, ranked, and re-merged. Reciprocal best-placed alignments are designated as “first pass”, while the non-best placed alignment is designated “second pass”.

