## Supplementary figures and images for "Interactive visualization of whole eukaryote genome alignments using NCBI’s Comparative Genome Viewer (CGV)"

### Supplemental Figure 1 (S1 Figure)

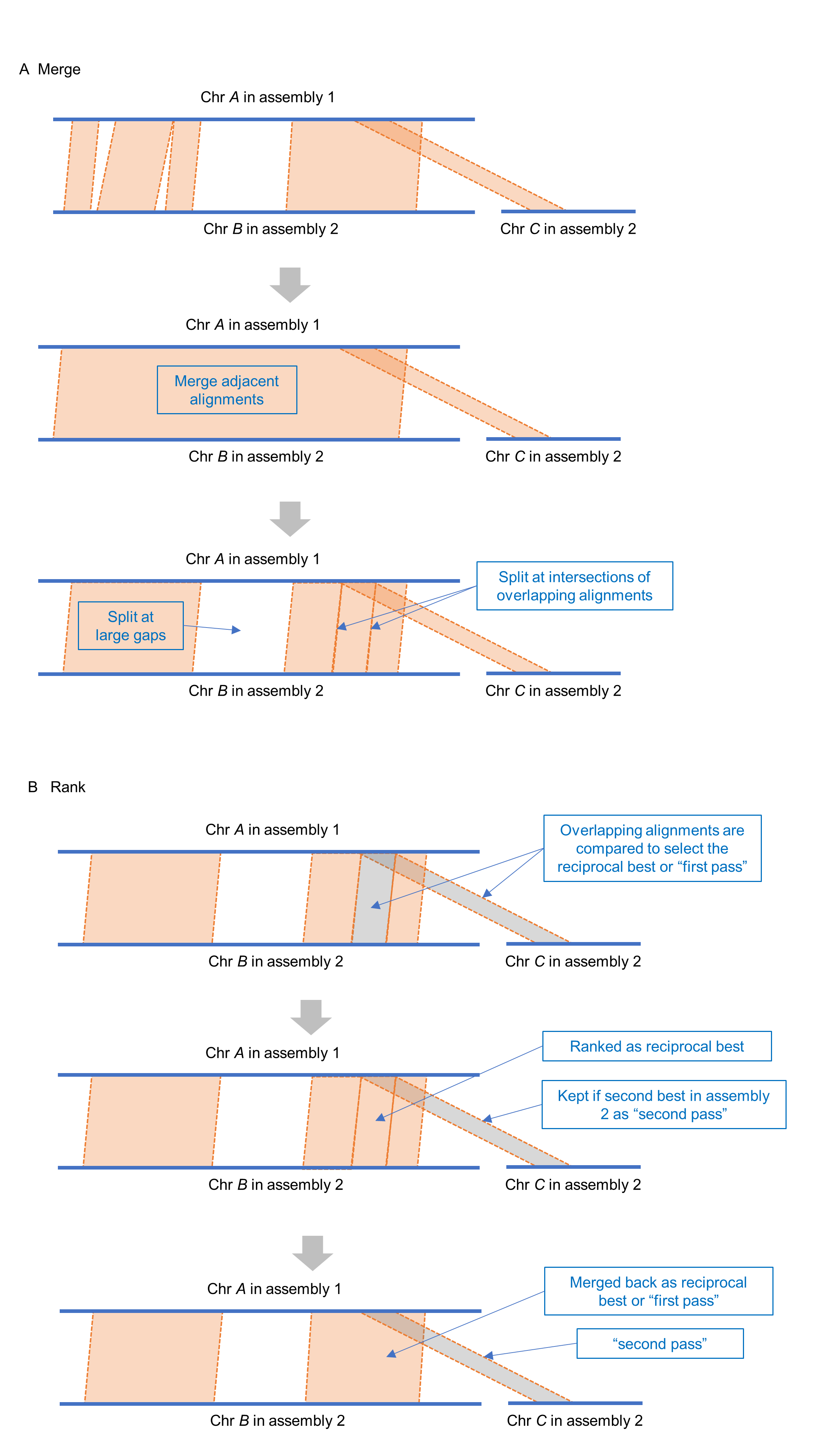
